# Lipidome unsaturation affects the morphology and proteome of the *Drosophila* eye

**DOI:** 10.1101/2023.05.07.539765

**Authors:** Mukesh Kumar, Canan Has, Khanh Lam-Kamath, Sophie Ayciriex, Deepshe Dewett, Mhamed Bashir, Clara Poupault, Kai Schuhmann, Oskar Knittelfelder, Bharath Kumar Raghuraman, Robert Ahrends, Jens Rister, Andrej Shevchenko

**Affiliations:** Max Planck Institute of Molecular Cell Biology and Genetics, Pfotenhauerstrasse 108, 01307 Dresden, Germany; Department of Biology, University of Massachusetts Boston, Integrated Sciences Complex, 100 Morrissey Boulevard, Boston, MA, USA

**Keywords:** *Drosophila*, retina, diet, vitamin A, fatty acid, proteome, lipidome, phototransduction

## Abstract

While the proteome of an organism is largely determined by the genome, the lipidome is shaped by a poorly understood interplay of environmental factors and metabolic processes. To gain insights into the underlying mechanisms, we analyzed the impacts of dietary lipid manipulations on the ocular proteome of *Drosophila melanogaster*. We manipulated the lipidome with synthetic food media that differed in the supplementation of an equal amount of saturated or polyunsaturated triacylglycerols. This allowed us to generate flies whose eyes had a highly contrasting length and unsaturation of glycerophospholipids, the major lipid class of biological membranes, while the abundance of other membrane lipid classes remained unchanged. By bioinformatically comparing the resulting ocular proteomic trends and contrasting them with the impacts of vitamin A deficiency, we identified ocular proteins whose abundances are differentially affected by lipid saturation and unsaturation. For instance, we unexpectedly identified a group of proteins that have muscle-related functions and increase their abundances in the eye upon lipidome unsaturation but are unaffected by lipidome saturation. Moreover, we identified two differentially lipid-responsive proteins involved in stress responses, Turandot A and Smg5, whose abundances decrease with lipid unsaturation. Lastly, we discovered that the ocular lipid class composition is robust to dietary changes and propose that this may be a general homeostatic feature of the organization of eukaryotic tissues, while the length and unsaturation of fatty acid moieties is more variable to compensate environmental challenges. We anticipate that these insights into the molecular responses of the *Drosophila* eye proteome to specific lipid manipulations will guide the genetic dissection of the mechanisms that maintain visual function when the eye is exposed to dietary challenges.

## Introduction

The proteome, the full complement of proteins that are produced by an organism, is largely determined by the genome. In contrast, the lipidome and the metabolome result from a complex and poorly understood interplay between environmental factors as well as developmental and metabolic processes. For instance, the composition of the lipidome is affected by dietary or temperature challenges that affect the length and unsaturation of fatty acid moieties in glycerophospholipids (GPL) ^1,2^, the major components of biological membranes. However, it remains unclear if the altered unsaturation of membrane lipids leads to marked changes in the proteome, for instance by affecting the abundances of lipid-binding proteins or membrane proteins. Here, we analyze the impacts of dietary lipid manipulations on the eye proteome of *Drosophila melanogaster*.

The *Drosophila melanogaster* eye is a membrane-rich organ that contains relatively low levels of storage lipids such as di- and triacylglycerols (DG and TG, respectively) but is enriched in GPLs ^3^. The GPLs exclusively comprise saturated or mono-unsaturated fatty acid moieties because flies lack Δ-6 / Δ-5 desaturases ^4^. However, others and we ^1,3,5^ have previously noticed that flies are able to incorporate a sizable proportion of dietary polyunsaturated fatty acids (PUFAs) into their compound eyes. PUFAs are critical for vision because dietary PUFA deficiency reduces the sensitivity of the photoreceptors, slows their photomechanical responses to visual stimuli ^6^, and impairs synaptic transmission ^7^. These visual defects are alleviated by supplementing the diet with specific PUFAs ^6,7^.

Since the composition and nutritional value of commonly used ‘standard’ laboratory foods can differ substantially, we previously developed a minimal and lipid depleted ‘M1’ food medium that we supplemented with the plant sterol stigmasterol and the vitamin A precursor beta-carotene ^8 9^. We showed that the morphology of the *Drosophila* compound eye and the light-sensing compartments of the photoreceptors (rhabdomeres) were wild type in flies reared on M1-food ^8^. Moreover, we recently studied the effects of vitamin A deficiency on the lipidome, proteome, and structure of the eye by omitting beta-carotene from M1-food (resulting in ‘M0-food’) ^8^. In the current study, we manipulated the lipidome by using M1-food as the basis to generate two food media that we supplemented with an equal amount of synthetic saturated (M3-food) or polyunsaturated triacylglycerols (M2-food), respectively. The M2- and M3-food media allowed us to raise flies whose eyes had a highly contrasting length and unsaturation of major GPLs, while the abundance of other membrane lipids (*e*.*g*., sphingolipids or sterols) remained unchanged. By bioinformatically comparing the ocular proteomic trends that resulted from rearing *Drosophila* on the four different but related diets (M0-M3 foods), we identified groups of proteins whose abundances are affected by lipid unsaturation.

## Results

### Experimental rationale and the design of synthetic food media for dietary interventions

We aimed to delineate how manipulations in lipid unsaturation affect the *Drosophila melanogaster* eye proteome. To this end, we altered the unsaturation of the eye lipidome by rearing flies on a set of four compositionally related food media (M0, M1, M2 and M3, **Table 1**).

**Table 1.** Overview of ‘M’ food composition

| Food | Yeast extract | Beta-carotene | Stigmasterol | TAG 42:0<br>TAG 48:0<br>TAG 54:0 | TAG 66:18 |
| --- | --- | --- | --- | --- | --- |
| M0<sup>(2)</sup> | x |  | x |  |  |
| M1 | x | x | x |  |  |
| M2<sup>(2)</sup> | x | x | x |  | x |
| M3 | x | x | x | x |  |

<sup>1</sup> added components are designated with 'x'.
<sup>2</sup> flies reared on these foods showed decreased rhabdomere size.

As their basis, we used M1-food, a soluble budding yeast extract that we supplemented with the vitamin A precursor beta-carotene ^9^ and the phytosterol stigmasterol ^8^. Except for the added stigmasterol, M1-food was lipid-depleted and thus an optimal starting composition for our specific lipid interventions. To provide different dietary fatty acids, we supplemented M1-food with an equal amount of unsaturated (M2-food; TAG 22:6/22:6/22:6) or saturated (M3-food; an equal mixture of TAG 14:0/14:0/14:0, TAG 16:0/16:0/16:0, and TAG 18:0/18:0/18:0) triacylglycerols (TAGs, **Table 1**). Effectively, M2-food supplied the omega-3 fatty acid DHA that flies metabolize to C20:5, C20:4, or even shorter PUFAs and use them for the synthesis of glycero- and glycerophospholipids ^4^. Conversely, M3-food closely resembled the common yeast diet since it supplied the flies with short, saturated fatty acids such as C14:0, C16:0, and C18:0. Consistent with this dietary similarity, saturated TAGs have been commonly detected in *Drosophila melanogaster* ^3,10^. In contrast, M2-food supplied unsaturated fatty acids that flies cannot generate *de novo*, but that can be metabolically derived from C22:6 ^4^ (see below). For clarity, we will hereafter refer to flies that were reared on the respective ‘M-’ food as ‘M-’ flies.

### Lipidome and proteome of M3-flies

We previously reported that the morphology of the compound eye and the light-sensing compartments of the photoreceptors (rhabdomeres) ^11^ of flies raised on M1-food closely resembled flies raised on ‘standard’ lab food (SF, **Materials and methods**) ^8^. Next, we used the M1-derived M2- and M3-foods (**Table 1**) to generate and compare flies with drastically different ocular lipid saturation. First, we tested if M3-food (M1-food supplemented with saturated TAGs that are common in *Drosophila*) affected the eye morphology, lipidome, and proteome (**Figure 1**). The M3 diet caused no obvious structural eye defects: the external compound eye morphology (**Figure 1A**), rhabdomere structure and arrangement (**Figure 1B**), as well as expression of the major rhodopsin Rh1 (**Figure 1B**) were very similar to M1- and SF-flies ^8^. Moreover, the rhabdomere cross-sectional area of the outer and inner photoreceptor types ^12^ was also not affected (p>0.05; **Figure 1C**).

**Figure 1:**
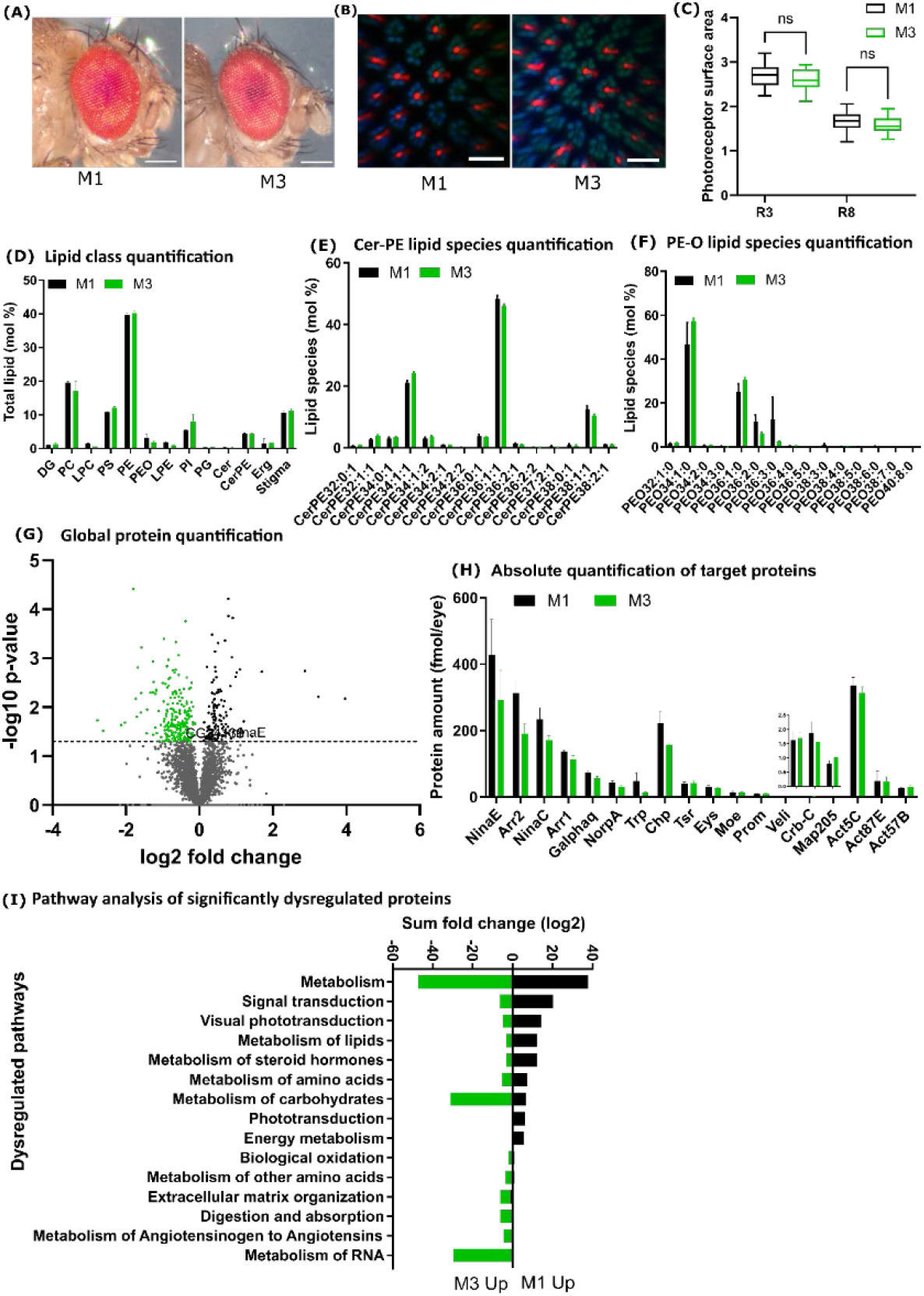
Effect of saturated lipids on structure, proteome, and lipidome of the *Drosophila* eye. **(A)** Compound eyes of wild-type male flies that were raised on M3-food (M1-food supplemented with saturated TAGs) or M1-food (control), respectively. **(B)** Confocal whole-mount images of male retinas; seven normally shaped rhabdomeres (Phalloidin, green) are visible for each unit eye. (**C**) Quantification of the cross-sectional areas of the rhabdomeres of two different photoreceptor types (outer: R3, inner: R8) for M3-food and M1-food. Data are for male retinas. (**D**) Eye lipidome analysis. Quantification (mol %) of different lipid classes in the *Drosophila* eye for M3- and M1-food. (**E**) Analysis of CerPE lipid species in the eye for M3- and M1-food. (**F**) Analysis of PE-O lipid species in the eye for M3- and M1-food. (**G**) Eye proteome analysis. The volcano plot shows differentially expressed proteins between M3 and M1 diet. (**H**) Quantification of the absolute (molar) abundances of proteins that play a major role in photoreceptor morphology or phototransduction for M3- or M1-food. (**I**) Significantly (p-value ≤ 0.05) dysregulated proteins from flies raised on M3- or M1-food were analyzed for GO term enrichment. Note that phototransduction was significantly downregulated.

Next, we subjected ten eyes of M1- and M3-flies, respectively, to shotgun lipidomics to quantify the molar abundance of 194 lipid species from eleven lipid classes, including the major membrane lipid classes LPE, PE, PE-*O*, Cer, Cer-PE, PC, LPC, PS, PI, DG, and PG. We found no significant difference in the mol% of lipid classes (**Figure 1D**) or lipid species composition of both sphingolipids (**Figure 1E**) and GPLs (**Figure 1F**). Taken together, supplementation of M1-food with saturated TAGs had no apparent impact on the major lipid classes in the eye.

A global eye proteome analysis by GeLC-MS/MS of M3-flies revealed that 221 proteins were significantly increased, and 120 proteins were significantly decreased in their abundances (**Figure 1G**) as compared to M1-flies. Subsequent pathway analysis revealed that the affected proteins were involved in ‘metabolism of proteins’ (15 proteins), ‘metabolism of carbohydrates’ (16 proteins), metabolism of lipids (five proteins), ‘metabolism of RNA’ (11 proteins), ‘signal transduction’ (11 proteins), and ‘phototransduction’ (six proteins). The significant alteration of metabolic pathways in the eye is consistent with the fact that due to the added TAGs, M3-food has a higher caloric value and also supplies flies with common fatty acids that can be directly used for lipid metabolism (**Figure 1 I**).

Since we detected no apparent changes in photoreceptor morphology or Rhodopsin expression, we next analyzed whether the saturated TAGs caused more subtle abundance changes in important photoreceptor proteins (**Figure 1 H**). To this end, we used our MS Western method ^13^ to determine the absolute (molar) abundances of 43 proteins that play key roles in phototransduction as well as the development and maintenance of photoreceptors ^8,14^. Notably, compared to M1-flies, the abundance of phototransduction proteins ^15^ decreased by 1.4-to 1.6-fold in M3-flies with no significant changes in their molar ratios. In contrast, the abundance of morphology-related proteins (*e*.*g*., Actins, Veli, Crb) was unchanged, consistent with the wild type morphology of the compound eyes and the photoreceptors.

We conclude that the dietary supply of short-chain saturated TAGs, which are compositionally similar to commonly found TAGs in *Drosophila* ^3,10^, did not significantly impact the morphology or the lipidome composition of the eye. However, adding the saturated TAGs to lipid-depleted M1-food altered several metabolic pathways related to protein, carbohydrate, lipid, and RNA metabolism, and also decreased the molar abundances of major phototransduction proteins.

### M2-flies show a perturbed photoreceptor morphology

To analyze the impact of increased unsaturation of the ocular lipidome, we reared flies on M2-food that contained a synthetic TAG with three moieties of docosahexaenoic acid (DHA, C22:6). While M2-flies had a normal external eye morphology (**Figure 2A**), their Rh1 staining was weaker and their rhabdomere cross-sectional area was significantly reduced compared to M3-flies (p < 0.001; **Figure 2B, C**) and M1-flies (see above) ^8^. Taken together, oversupplying a polyunsaturated TAG decreased Rh1 expression and rhabdomere size in M2-flies.

**Figure 2:**
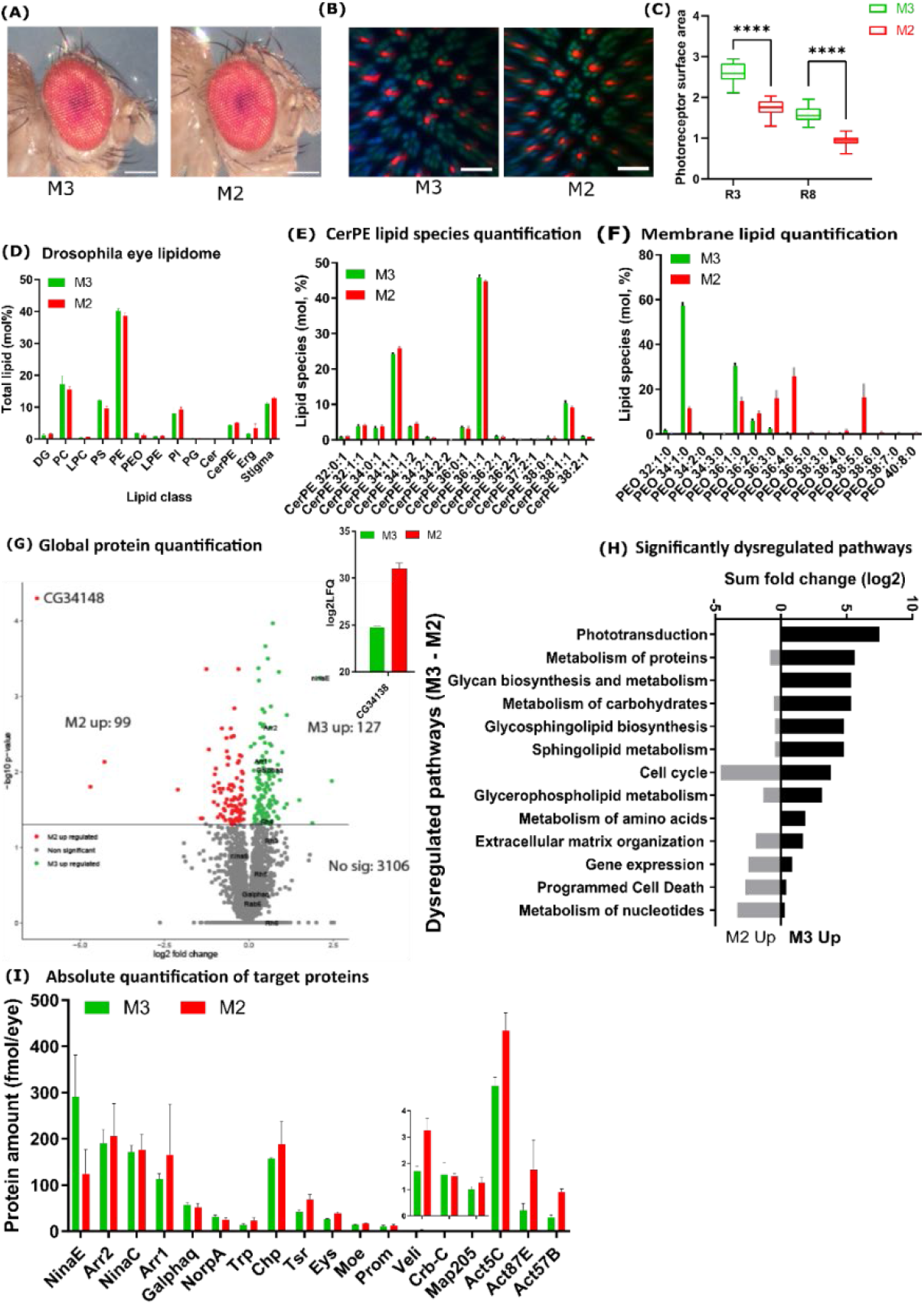
Effects of highly saturated and unsaturated lipids on the structure, proteome, and lipidome of the *Drosophila* eye. **(A)** Compound eyes of wild-type male flies that were raised on M2- or M3-diet, respectively. **(B)** Confocal whole-mount images of male retinas; seven rhabdomeres are visible (Phalloidin, green) for each unit eye. Note the reduced rhabdomere diameter in the case of M2 diet. (**C**) Quantification of the cross-sectional areas of the rhabdomeres of two different photoreceptor types (outer: R3, inner: R8) for M2 and M3 (control) diet. (**D**) Eye lipidome analysis. Quantification (mol %) of different lipid classes in the *Drosophila* eye for M2 and M3 diet. (**E**) Analysis of CerPE lipid species in the eye for M2 and M3 diet. (**F**) Analysis of PE-O lipid species in the eye for M2 and M3 diet. (**G**) Eye proteome analysis. The volcano plot shows differentially expressed proteins between M2 and M3 diet. The inset shows the quantification of the protein CG34138, which is highly upregulated in response to increased lipid unsaturation. (**H**) Significantly (p-value ≤ 0.05) dysregulated proteins from flies raised on M2 or M3 diet were analyzed for GO term enrichment. Note that phototransduction was significantly downregulated in the case of M2 diet. (**I**) Quantification of the absolute (molar) abundances of proteins that play a major role in photoreceptor morphology or phototransduction for M2 and M3 diet.

### Eye lipidome of M2-flies is highly unsaturated

Despite the major difference in unsaturation between the supplemented dietary TAGs, the ocular lipid class composition of M1-^8^, M2-, and M3-flies was surprisingly similar (**Figure 2D**). However, the molecular species profile in M2-flies changed in a lipid-class dependent manner: GPLs incorporated a variety of PUFAs that were metabolically derived from the DHA of the supplied unsaturated TAG. We detected unsaturated species with up to six double bonds in all major GPL classes (PC, PE, PS, PI, PG) but also in ether PEs (PE-*O*) ^3^ (**Figure 2D, F**). Furthermore, the absolute abundance of shorter and more saturated lipids (zero to two double bonds per lipid molecule) was reduced. In contrast, the molecular profiles of *lyso-*PC and *lyso-*PE, but also of major energy storage lipids such as diacylglycerols (DG), were largely unaffected (**Figure 2D**). This suggests that the metabolic conversion of DHA to C20:5, C20:4, and down to C18:2 could be a rate-limiting step. Lastly, supplying DHA did not increase the production of sphingolipids with elongated and/or unsaturated N-acylamidated fatty acid moieties. Altogether, the ocular membrane lipidome of M2-flies became markedly unsaturated.

### Proteome-wide impact of lipidome unsaturation

The M2-and M3-foods share the same compositional basis, M1-food (**Table 1**), and provide flies with the same amount of dietary TAGs. Yet, these TAGs differ in the unsaturation of their fatty acid moieties, which resulted in the markedly higher unsaturation of GPLs -major constituent of lipid membranes. At the same time, other membrane lipids – sphingolipids and stigmasterol – remained unchanged. To assess the proteome-wide impact of membrane lipidome unsaturation, we compared the ocular proteomes of M2- and M3-flies. Out of the total of 3106 quantified proteins, 99 were significantly upregulated and 127 downregulated in M2-flies. In both M2-*vs*. M3-flies (**Figure 1H and Figure 2H**) and M1-*vs*. M3-flies, the protein changes were mostly associated with altered metabolism and affected similar metabolic pathways, albeit with a different magnitude. In contrast to M1- and M3-flies, both the lipidome and the rhabdomere morphology were affected in M2-flies (**Figure 2B, C**). Conversely, vitamin A-deprivation by M0-food (i.e., M1-food lacking the vitamin A precursor beta-carotene) (**Table 1**) also affected rhabdomere morphology, but not the eye lipidome ^8^. Taking the ocular impacts of all four diets (M0-M3) into account, we hypothesize that the eye proteome composition were influenced by three main factors (summarized in **Table 2):** *i)* rhabdomere morphology defects; *ii)* global changes in metabolism due to the added TAGs, irrespective of their fatty acid moieties; and *iii)* enhanced unsaturation of membrane lipids caused by incorporation of PUFAs that were derived from the highly unsaturated dietary TAG.

**Table 2.** Rationale behind successive subtraction of proteomic datasets^1^

| Compared proteomes<sup>2</sup> | Reflects impact of: |  |  |
| --- | --- | --- | --- |
|  | supplied TAGs | decreased rhabdomere size | unsaturated GPLs |
| M3 vs. M2 | x | x | x |
| M1 vs. M0 |  | x |  |
| M3 vs. M1 | x |  |  |

<sup>1</sup>The anticipated impact of factors revealed by the pairwise comparison is marked with 'x'.
<sup>2</sup>Comparison of the proteomes of corresponding M-flies

We further hypothesize that the proteome changes in M2-flies (as compared to M1-flies) overlap with those of M0-flies because both dietary manipulations decreased the rhabdomere size. Lastly, we also expect an overlap between the proteome changes in M2- and M3-flies due to a general metabolic response to a lipid-rich diet. Hence, we took advantage of these four (M0-, M1-, M2- and M3-flies) proteomic datasets to perform a pairwise comparison and successive subtraction of similarly regulated proteins in multiple datasets to identify those, whose abundance specifically responds to membrane lipid unsaturation.

### Identification of proteins that respond to lipidome unsaturation

To identify ocular proteins that respond to lipidome unsaturation, we employed the following data processing workflow: we filtered the datasets with two permissive cut-off thresholds for p-value (*p*<0.05 or -log10 > 1.3) and fold change (FC; |FC| >1.5 or |log2 FC| >0.58), where +/-indicates the direction of change in the M-food comparison. For clarity, we assigned a negative FC to proteins enriched in M*y* in each M*x vs*. M*y* comparison. Next, from the list of significantly changed proteins in the M3-flies *vs*. M2-flies comparison (hereafter termed [M3 *vs*. M2]), we subtracted the list [M1 *vs*. M0], which yielded the {[M3 *vs*. M2] - [M1 *vs*. M0]} list. We chose to disregard |FC| values and only subtracted proteins whose abundance changed in the same direction in both comparisons. For example, a protein was removed from the [M3 *vs*. M2] list if it was upregulated in M2-flies, but also in M0-flies in the [M1 *vs*. M0] comparison, irrespective of the FC magnitude. In this way, we removed proteins from the [M3 *vs*. M2] list, whose change in abundance was associated with abnormal rhabdomere morphology and not a specific response to unsaturation (**Table 2**). We applied the same criteria to further subtract proteins that responded similarly to adding TAGs ([M3 *vs*. M1]), i.e., that were not differentially affected by the different TAGs.

Altogether, the final list {[M3 *vs*. M2] - [M1 *vs*. M0] - [M1 *vs*. M3]} comprised twenty-five proteins that are associated with ocular lipidome unsaturation, and five of these proteins were membrane-associated. In comparison, out of the total of 3106 proteins quantified in M2-flies, 910 proteins were membrane-associated. Strikingly, the higher unsaturation of membrane lipids therefore had no impact on the abundance of membrane proteins *en masse*.

#### Abundance increase with lipidome unsaturation, unaffected by lipid saturation

a group of seven proteins was upregulated in M2-flies (unsaturated), but unaffected in M0-flies (vitamin A deprived) and M3-flies (saturated). Unexpectedly, a GO term analysis revealed that the affected proteins all have muscle-related functions and include: Myofilin (a structural component of the thick muscular filaments) ^16^, Paxillin (a cytoskeletal adaptor protein that regulates cell fusion in muscles) ^17^, Paramyosin (an invertebrate-specific muscle protein that is part of the thick filament), Myosin heavy chain, Upheld (encodes the calcium-binding muscle regulatory protein Troponin T) ^18^, Wings up A (a cytoskeletal protein of the Troponin complex), and RIM-binding protein (an active zone protein involved in neuromuscular synaptic transmission).

#### Abundance decrease with lipidome unsaturation, increase with saturation

Conversely, our analysis identified a group of six proteins whose abundance decreased in ‘unsaturated’ M2-flies and increased in ‘saturated’ M3-flies. This group includes the putative TAG lipase CG5162, whose human orthologs are implicated in hyperlipidemia and obesity, and two lipid transporter proteins: the scaffolding apolipoprotein Apolipophorin is a member of the conserved ApoB family and is involved in the synthesis of the lipoprotein lipophorin, which transports lipids between tissues. Crossveinless d is another lipoprotein that resembles vitellogenins, a major component of embryonic yolk ^19^. Notably, this group also included two proteins that mediate stress responses, Turandot A and Smg5. The humoral factor Turandot A (TotA) exhibited the strongest (over 4-fold) downregulation among all proteins in response to lipidome unsaturation and has been shown to respond to various types of environmental stresses ^20^. Smg5 is an essential nonsense-mediated mRNA decay factor that was found in an obesity screen to regulate TAG levels specifically in muscle cells ^21,22^. Lastly, the two predicted plasma membrane metallopeptidases angiotensin-converting enzyme Ance-4 and Neprilysin 6 were both downregulated in M2-flies, albeit Neprilysin 6 was not significantly changed in M3-flies ^23,24^.

#### Abundance increase with lipidome unsaturation, decrease with lipidome saturation

Interestingly, the 130 kDa Golgi matrix protein GM130 showed the reverse lipid response pattern: it was downregulated in the eyes of M3-flies, but upregulated in M2-flies. GM130 is a structural protein that is involved in connecting the Golgi compartments in the soma and dendrites of neurons ^25^.

Finally, we identified several proteins that differentially responded to the two lipid manipulations and have important roles in visual signaling. Two of these proteins are involved in visual pigment synthesis and show an abundance increase upon lipid unsaturation: the chaperone NinaA is required for the synthesis of the visual pigments Rh1 and Rh2 ^26^ and the oxidoreductase NinaG is essential for the synthesis for the vitamin A-derived chromophore ^27,28^. Conversely, the protein levels of the light-sensitive cation channel Trpl and the eye-specific protein kinase InaC ^15^ both decreased upon lipid unsaturation.

In addition to the global proteome analysis, we again quantified the changes in absolute (moles per eye) protein abundances with our targeted MS Western method ^13^ (**Figure 2I**). In the eyes of M2-flies, the molar abundance of phototransduction proteins was significantly lower compared M1-flies, which was very similar to (and not significantly different from) M3-flies. However, Rh1 was significantly reduced compared to M3-flies. Conversely, the abundance of structural proteins (*e*.*g*., Actins and Veli) increased by approx. 2.2x, consistent with the increase of other proteins of the actomyosin machinery.

Taken together, we analyzed the effects of lipidome manipulations on the ocular proteome in four related synthetic diets. We observed that a dietary manipulation that caused an increased unsaturation of the eye lipidome elicited a specific proteome response that differentially changed the abundances of proteins involved in lipid metabolism and transport, muscle organization, stress responses, and visual signaling.

## Discussion

### Muscle-related proteins respond to lipid unsaturation in the eye

We studied the effects of lipidome manipulations on the ocular proteome and need to emphasize that grouping differentially expressed proteins by their lipid response patterns does not necessarily imply a similar function or molecular relationship. Yet, we surprisingly identified a group of seven proteins that have muscle-related functions and increase their abundances in the eye upon lipidome unsaturation (but are unaffected by lipidome saturation). This result can be interpreted in several ways: the differential expression of these muscle-related proteins could reflect a reorganization of muscle tissue in the eye in response to increased lipid unsaturation. For instance, this could involve the ocular muscles that move the *Drosophila* retina to track motion and stabilize the retinal image ^29^. Furthermore, the actomyosin machinery is required for the formation of the luminal matrix space between the rhabdomeres ^30^ and actomyosin contraction plays a critical role in shaping the morphology of the eye ^31^. Alternatively, it is conceivable that these proteins have other, yet to be identified functions in the *Drosophila* eye that are required upon increased lipid unsaturation. For instance, the expression of genes that are associated with muscle development has been detected in the pupal eye ^32^ and Troponin I/Wings up A additionally controls the proliferation of epithelial cells as well as the localization of apico-basal polarity signaling proteins ^33^. Third, the upregulation of muscle-related proteins could be due to a yet to be identified mechanistic link between lipid unsaturation and increased muscle protein expression that co-occur at low temperatures: we previously showed that at temperatures below 15°C, *Drosophila* alter their dietary preference from yeast to plant material, which provides unsaturated fatty acids that improves membrane fluidity and motor functions ^1^. Notably, another study revealed that the expression of genes that encode the myosin heavy and light chains is upregulated at low temperatures in adult *Drosophila*, potentially as an adaptive response to compensate for decreased muscle contractility and to maintain flight performance at low temperatures ^34^. These data suggest that the muscle machinery is plastic and can adapt to both dietary and temperature stresses.

### Stress-responsive proteins are downregulated upon lipid unsaturation

We also identified two differentially lipid-responsive proteins involved in stress responses, Turandot A and Smg5, whose abundances decrease with lipid unsaturation. Smg5 is an essential nonsense-mediated mRNA decay factor that was found in an obesity screen to regulate TAG levels specifically in muscle cells ^21,22^. The humoral factor Turandot A (TotA) showed the strongest (over 4-fold) downregulation among all proteins in response to lipidome unsaturation. *TotA* expression is induced by various environmental stresses such as UV light, heat, cold ^35^, bacterial infection, and oxidative agents ^20^. Moreover, *TotA* expression is also increased when flies adapt to a high-protein-low-carbohydrate diet ^36^. It is unclear how a broad range of stress stimuli increases *TotA* transcription, while lipid unsaturation decreases the TotA protein levels. Yet, in another study, a high-fat diet caused a *TotA* downregulation specifically in males ^37^. Since TotA is regulated by JAK-STAT and MAPK pathways ^38^, its downregulation could reflect a lower activity of those pathways or an overall reduction in stress load ^39^.

### Robustness of ocular lipid class composition and membrane protein expression

*Drosophila* lacks the ability to synthesize long-chain PUFAs from shorter-chain precursors (such as linoleic acid, C18:2) and the rhabdomere membranes lack lipids with PUFA moieties of more than 18 carbon atoms ^4,10^. The TAG that we added in the case of M2 food supplied the omega-3 PUFA DHA, which has not been detected in *Drosophila* ^4,40^, but is present at high levels in the rhabdomere-equivalent outer segments of mammalian photoreceptors ^41-43^. Unexpectedly, we discovered that once the supplied DHA was metabolized to generate various long-chain polyunsaturated GPLs, the lipid class composition of the eye remained unchanged. Moreover, DHA did not affect the molecular profile of major sphingolipids (Cer and Cer-PE). Others ^44^ and we ^1,3,5^ previously observed similar homeostatic trends when flies were switched from a ‘mostly saturated’ yeast diet to ‘mostly unsaturated’ plant diets. Therefore, we speculate that this robustness of the ocular lipid class composition to dietary changes may be a general homeostatic feature of the organization of eukaryotic tissues, while the length and unsaturation of fatty acid moieties is more variable, potentially to allow a compensatory response towards environmental and dietary challenges ^1,2^. Another unexpected observation was that the increased unsaturation of membrane lipids in the fly eye affected the abundance of only very few membrane proteins. This robustness of the expression of membrane proteins also suggests that, once formed, lipid-protein assemblies can be incorporated into membranes of variable composition and properties.

### Effects of dietary manipulations on photoreceptor morphology and phototransduction protein expression

Photoreceptors have highly specialized light-sensing compartments that are called rhabdomeres in flies and outer segments in mammals ^11^. Altering the molecular composition of the rhabdomere membrane can affect its physical properties such as stiffness or fluidity ^45,46^ and thereby visual signaling ^6^: a PUFA-deficient yeast diet impairs the speed and sensitivity of the phototransduction cascade, potentially due to decreased rhabdomere membrane fluidity ^6^. In a previous study, we discovered that the expression of the components of the rhabdomeric phototransduction machinery is dependent on vitamin A ^8^; in the current study, we found that it is also dependent on lipid unsaturation. The impacts of M2-diet resemble vitamin A deficiency (M0-diet) because both cause a severely reduced rhabdomere size and decreased levels of phototransduction proteins ^8,9^. The M3-diet containing three short chain saturated TAGs also decreased the phototransduction protein levels but did not cause any obvious rhabdomere damage. This suggests that rhabdomere defects are not the cause for the decreased phototransduction protein expression, which is also consistent with our previous finding that *crumbs* mutants have severe rhabdomere defects, but do not exhibit significant changes in the levels of phototransduction proteins or Rh1 ^14^. The molecular mechanisms that underlie the impacts of these three different dietary manipulations on the expression levels of phototransduction proteins remain to be elucidated.

In conclusion, we anticipate that these insights into the molecular responses of the *Drosophila* eye proteome to specific lipid manipulations and the datasets that we generated will be useful resources for the genetic dissection of the mechanisms that maintain visual function when the eye is exposed to dietary challenges.

## Material and Methods

### *Drosophila* culture

The *Drosophila melanogaster* wild-type strains Oregon R and Canton S were reared under a 12h light/12h dark cycle at 25°C. The flies were raised on one of the five different food media (SF diet or M0-M3 diet) from the embryonic to the adult stage. Three-to-four days old male flies were collected and used for the experiments.

### *Drosophila* food media

*Drosophila* flies were raised on one of five different diets: ‘standard’ lab food (SF) or one of four synthetic media (M0-M3). SF contained per liter: 8g agar, 18g brewer’s yeast, 10g soybean, 22g molasses, 80g cornmeal, 80g malt, 6.3ml propionic acid, and 1.5g Nipagin. Minimal M0 food contained per liter: 10g UltraPure Agarose (Invitrogen), 100g yeast extract (Kerry), 100g glucose (Merck), 1.5mL Nipagin (Sigma-Aldrich) in 10% in ethanol, and 1 g stigmasterol (Sigma). M1 food is M0 food supplemented with 0.5g β-carotene (Sigma). M2 food is M1 food supplemented with an unsaturated TAG (TAG 66:18; Larodan) and M3 is M1 food supplemented with equal amounts of three saturated TAGs (TAG 42:0; TAG 48:0; TAG 54:0; Larodan).

### Imaging the *Drosophila* eye

The flies were immobilized by embedding them in an agarose gel. To prepare the agarose gel, we dissolved two grams of Ultra-pure Agarose (Invitrogen) in 100 mL distilled water. The solution was heated in a 500 mL Erlenmeyer flask for several minutes until bubbles were visible. The flask was then placed in a water bath (Thermo Scientific™) set to 58° C to cool the solution down to 58° C. Next, flies were anesthetized with carbon dioxide gas and transferred to a 60 mm petri dish (Falcon®) filled with about 10 mL of the liquid agarose gel. We then submerged the wings and legs using forceps and placed the petri dish on ice for the gel to solidify. After solidification of the gel, the petri dish was placed under the Stemi 508 Trinoc microscope (Zeiss model #4350649030), and the fly head was adjusted with the forceps such that one eye was oriented in parallel to the microscope lens. The petri dish was placed back on ice until imaging. Imaging was performed with an Axiocam 208 HD/4k color camera (model #4265709000), which was set to auto exposure and auto white balance. Pictures were processed with Fiji, Adobe Photoshop 2020, and Adobe Illustrator 2020 software.

### Confocal microscopy and immunohistochemistry of *Drosophila* photoreceptors

We dissected retinas of four days old flies as previously described ^47^ in cold PBS. We fixed the retinas in 3.8% formaldehyde solution before removing the laminas as well as head cuticle. We then incubated the retinas overnight in the primary antibody (mouse anti-Rh1 4C5, 1:10, from Developmental Studies Hybridoma Bank, University of Iowa) that was diluted in PBST (PBS + 0.3% Triton-X, Sigma). We removed the primary antibody solution the next morning and washed the retinas three times with PBST. In the evening, we incubated the retinas in secondary antibody diluted in PBST (1:800, Alexa Fluor 647-conjugated raised in donkey, Invitrogen) and Alexa Fluor 488-conjugated Phalloidin (1:100, Invitrogen). We performed three PBST washes next morning. Next, we mounted the retinas using SlowFade (Molecular Probes) on a bridge slide and imaged them with a Zeiss LSM 8 confocal microscope. We processed the raw images with Fiji (https://imagej.net/software/fiji/) and further cropped and contrasted them with Adobe Photoshop and Adobe Illustrator.

### Rhabdomere measurements and statistics

We used Fiji’s measurement tool on the Phalloidin channel (green) to quantify the cross-sectional area of the rhabdomeres in eight unit eyes from five different retinas. First, we used the freehand selection tool to draw a circle around the brightest part of one of the R3 rhabdomeres at the level of the R8 rhabdomeres. Second, we used the ROI manager tool to measure the area of the circled rhabdomere. Third, we used RStudio (https://www.rstudio.com/) to perform an ANOVA followed by a Tukey honestly significant difference (Tukey’s HSD) test to determine whether the size differences were statistically significant between experimental (i.e. food media) groups. The significance cut-off was p<0.05; we then generated Box-and-Whisker plots in RStudio.

### Lipid extraction and shotgun lipidomics analysis of the *Drosophila* eye

Whole eyes (n=10) were dissected with a thin blade and placed in 40 µL of 150 mM ammonium bicarbonate buffer containing 10% of isopropanol (IPA) into a 2 mL Eppendorf tube and snap-frozen in liquid nitrogen and stored at -80°C or processed immediately. The eyes were mechanically disrupted with 1 mm zirconia beads, samples were dried under vacuum to remove isopropanol. The total lipids were extracted using the methyl tert-butyl ether (MTBE) extraction according to Matyash et al., 2008. Samples were resuspended in 200 µL of water and 700 µL of MTBE/methanol (5:1.5, v/v) containing internal standards (0.539 nmol zymosterol-d5, 0.782 nmol stigmasterol-d6, 0.313 nmol triacylglycerol-d5 50:0, 0.073 nmol diacylglycerol-d5 34:0, 0.138 nmol PC 12:0/13:0, 0.109 nmol LPC 13:0, 0.067 nmol PS 12:0/13:0, 0.147 nmol PE 12:0/13:0, 0.053 nmol LPE 13:0, 0.090 nmol PI 12:0/13:0, 0.068 nmol PG 12:0/13:0, 0.102 nmol Cer 30:1, 0.077 nmol PA 12:0/13:0, 0.068 nmol GalCer 30:1, 0.081 nmol LacCer 30:1, 0.074 nmol CerPE 29:1). After centrifugation, the organic phase was collected and dried under vacuum to avoid lipid oxidation. The whole extraction procedure including sample preparation was performed at 4°C in order to prevent lipid degradation. Mass spectrometric analyses were performed on a Q Exactive instrument (Thermo Fisher Scientific, Bremen) equipped with a robotic nanoflow ion source TriVersa NanoMate (Advion BioSciences, Ithaca, NY) using chips with the diameter of spraying nozzles of 4.1 mm. Lipids were identified by LipidXplorer software (Herzog et al, 2011) by matching m/z of their monoisotopic peaks to the corresponding elemental composition constraints.

### Protein extraction and GeLC-MS/MS analysis of the *Drosophila* eye proteome

The compound eyes (n=40) were dissected from the male flies raised under different food conditions (see above) and placed in lysis buffer containing 150 nM NaCl, 1 mM EDTA, 50 mM Tris-HCl (pH7.5), 1 tablet Roche protease inhibitors, 0.2% w/v CHAPS, 0.1% w/v OGP, 0.7% v/v triton X-100, 0.25 μg/mL DNase and RNase. The samples were immediately snap frozen using liquid nitrogen and stored at -80°C or immediately processed. The eye tissues were homogenized and to the supernatant an equal volume of 2X SDS Laemmli sample buffer (SERVA Electrophoresis GmbH, Heidelberg, Germany) was added. The samples were heated at 80°C for 10-15 minutes and were loaded on 4-20% 1D SDS PAGE. The protein bands were visualized by Coomassie Brilliant Blue staining. Each gel lane was cut into six gel slices, and each gel slice was co-digested with heavy isotope labeled CP02 and a gel band containing 1pmol of BSA standard.

### GeLC-MS/MS

In-gel digestion was carried out as previously described ^8^. Briefly, the electrophoresed gel rinsed with water was stained with Coomassie Brilliant Blue R-250 for 10 minutes at RT and then destained with destaining solution (Water: Methanol: Acetic acid, 50:40:10 (v/v/v). The gel slice was excised as per the expected molecular weight of the proteins of interest and further cut into small pieces (∼1 mm size). The gel pieces were then transferred into 1.5 ml LoBind Eppendorf tubes and further processed. The gel pieces were completely destained by ACN/Water, reduction was done by incubating the gel pieces with 10 mM Dithiothreitol at 56°C for 45 minutes. Alkylation was carried out with 55mM Iodoacetamide for 30 minutes in the dark at RT. The reduced and alkylated gel pieces were washed with water/ACN and finally shrunken with ACN, ice-cold trypsin (10ng/µl) was added to cover the shrunken gel pieces and after 1hr of incubation on ice, excess trypsin (if any) was discarded. The gel pieces were then covered with 10mM NH4HCO3 and incubated for 12-15hrs at 37°C. The tryptic peptides were extracted using water/ACN/FA, dried using a vacuum centrifuge and stored at -20°C until next use. The tryptic peptides were recovered in 5% aqueous FA and 5 μL were injected using an autosampler into a Dionex Ultimate 3000 nano-HPLC system, equipped with a 300 μm i.d. × 5 mm trap column and a 75 μm × 15 cm Acclaim PepMap100 C18 separation column. 0.1% FA in water and ACN were used as solvent A and B, respectively. The samples were loaded on the trap column for 5 min with a solvent A flow of 20 µL/min. The trap column was then switched online to the separation column, and the flow rate was set to 200nL/min. The peptides were fractionated using a 180 min elution program: a linear gradient of 0% to 30% B delivered in 145 min and then B% was increased to 100% within 10 min and maintained for another 5 min, dropped to 0% in 10 min and maintained for another 10 min. Mass spectra were acquired using either LTQ Orbitrap Velos or Q Exactive HF mass spectrometer both from Thermo Fisher Scientific (Bremen, Germany).

### Data processing for protein identification and quantification

Mascot v2.2.04 (Matrix Science, London, UK) was used for peptide identifications against the custom-made database containing the sequence of the target protein, to which sequences of human keratins and porcine trypsin were added. For eye proteome analysis, the *Drosophila* reference proteome database from UniProt was used. The database searches were performed with the following mascot settings: precursor mass tolerance of 5 ppm; fragment mass tolerance of 0.6 Da and 0.03 Da for the Velos and QE-HF data respectively; fixed modification: carbamidomethyl (C); variable modifications: acetyl (protein N-terminus), oxidation (M); Label: 13C (6) (K), Label: 13C (6) 15N (4) (R), 2 missed cleavages were allowed. More details of mascot search parameters are provided (Table 11). Progenesis LC-MS v4.1 (Nonlinear dynamics, UK) was used for the peptide feature extraction and the raw abundance of identified peptide was used for absolute quantification. MaxQuant v1.5.5.1 and Perseus v1.5.5.3 was used for label-free quantification and subsequent statistical analysis. MaxQuant analysis was done with default settings.

## Acknowledgements/funding

This publication was supported by the National Eye Institute of the National Institutes of Health under Award Number R01EY029659 to J.R. and internal funding of MPI CBG, Dresden. The content is solely the responsibility of the authors and does not necessarily represent the official views of the National Institutes of Health. The funders had no role in study design, data collection and analysis, decision to publish, or preparation of the manuscript.

